## Supplemental Information for "Neural network aided approximation and parameter inference of stochastic models of gene expression"

Jiang *et. al.*

### Supplementary Note 1 Exact solution for Model I: constitutive transcription with delayed degradation

Here we present the exact time-dependent solution of Model I which in the main text is used to illustrate the key ideas behind the artificial neural network (ANN) aided model approximation and also to evaluate its precision. Note that a more general derivation was reported in [1]. The model has two reactions:

$$\varnothing \xrightarrow{\rho} N, \quad N \underset{\tau}{\Rightarrow} \varnothing, \tag{S1}$$

where the first reaction describes the production of nascent RNA as a Poisson process with its mean equal to $\rho$, while the second reaction describes the "decay" of the nascent RNA a time $\tau$ later after it is produced (actually it models its detachment from the gene locus). The objective is to find the probability $P(n,t)$ that there are $n$ nascent RNAs at time $t$.

#### 1.1 Derivation of delay master equation

We next derive the delay CME describing the stochastic dynamics. We want to obtain the probability $P(n, t + \Delta t)$. There are three sets of events that contribute to this: (A) there are $n-1$ molecules at time $t$ and a nascent RNA molecule is produced in the time interval $(t, t+\Delta t)$; (B) there are $n+1$ molecules at time $t$ and a nascent RNA molecule is removed in the time $(t, t+\Delta t)$; (C) there are $n$ molecules at time $t$ and no reaction occurs in the time interval $(t, t+\Delta t)$.

Now in the infinitesimal limit $\Delta t \to 0$, the probability that event (A) occurs is simply the conventional propensity associated with a firing of the production reaction:

$$P(n, t+\Delta t; n-1, t) = \rho \Delta t P(n-1, t). \tag{S2}$$

The probability that event (B) occurs requires a careful consideration of the history of the process: (i) $(n', t-\tau)$ leads to $(n'+1, t-\tau+\Delta t)$; (ii) $(n'+1, t-\tau+\Delta t)$ leads to $(n+1, t)$; (iii) $(n+1, t)$ leads to $(n, t+\Delta t)$, where $n'$ is the number of nascent RNAs at time $t-\tau$. Intuitively since degradation is a delayed reaction, a removal reaction in (iii) is due to a production reaction in (i) a time interval $\tau$ earlier. The probability of (i) occurring is $\rho\Delta t P(n', t-\tau)$. The probability of (iii) occurring is 1 since every production event is followed by a removal event a time $\tau$ later. The probability of (ii) occurring is:

$$\begin{aligned} P(n+1, t | n'+1, t-\tau+\Delta t) &= P(n, t | n', t-\tau+\Delta t) \\ &= P(n, t | 0, t-\tau+\Delta t). \end{aligned} \tag{S3}$$

The two "+1"s in the left-hand-side of Eq. (S3) represent that a nascent RNA was born in time $(t-\tau, t-\tau+\Delta t)$ and stays in the system till time $t$, and notably does not participate in any reaction in time $(t-\tau+\Delta t, t)$; hence it follows that we can write the first line on the right hand side of Eq. (S3). Now since all nascent RNAs born before time $t-\tau$ (whose number is $n'$) are removed from the system, the probability $P(n, t|n', t-\tau+\Delta t)$ must be independent of $n'$ and the $n$ nascent RNAs at time $t$ are produced between time $(t-\tau+\Delta t, t)$; hence it follows that we can write the second line on the right hand side of Eq. (S3). The probability that event (B) occurs is given by the product of the probability of events (i), (ii) and (iii) and summing over all values of $n'$:

$$P(n, t+\Delta t; n+1, t) = \rho\Delta t \sum_{n'} P(n', t-\tau) P(n, t|0, t-\tau+\Delta t) = \rho \Delta t P(n, t|0, t-\tau+\Delta t), \tag{S4}$$

where $P(n, t|0, t-\tau+\Delta t)$ is the probability of having $n$ nascent RNAs produced in the time interval $(t-\tau+\Delta t, t)$.

See Fig. 1 for an illustration of the aforementioned considerations for event (B). The probability that event (C) occurs is obtained by subtracting all the probabilities of one reaction occurring from the probability $P(n,t)$.

From the law of total probability, we have:

$$P(n,t+\Delta t) = P(n,t+\Delta t; n-1,t) + P(n,t+\Delta t; n+1,t) \\ + [P(n,t) - P(n+1,t+\Delta t; n,t) - P(n-1,t+\Delta t; n,t)], \quad \text{(S5)}$$

which by means of Eqs. (S2) and (S4) simplifies to:

$$\begin{aligned} P(n,t+\Delta t) =& \rho\Delta t P(n-1,t) + \rho\Delta t P(n,t|0,t-\tau+\Delta t) \\ &+ P(n,t) - \rho\Delta t P(n,t) - \rho\Delta t P(n-1,t|0,t-\tau+\Delta t) \\ =& P(n,t) + \rho\Delta t[P(n-1,t) - P(n,t)] + \rho\Delta t[P(n,t|0,t-\tau+\Delta t) - P(n-1,t|0,t-\tau+\Delta t)] \end{aligned} \quad \text{(S6)}$$

Dividing $\Delta t$ on the both sides of Eq. (S6) and letting $\Delta t \to 0$, the delay CME of Model I is obtained:

$$\frac{\mathrm{d}P(n,t)}{\mathrm{d}t} = \rho[P(n-1,t) - P(n,t)] + \rho[P(n,t|0,t-\tau) - P(n-1,t|0,t-\tau)]. \quad \text{(S7)}$$

#### 1.2 Solution of the delay master equation

To solve Eq. (S7), one has to find an equation governing the conditional probability $P(n,t|0,t-\tau)$. Now we know that the $n$ nascent RNAs produced during time $(t-\tau,t)$ and any nascent RNA produced prior to $t-\tau$ cannot survive up to time $t$. In other words, the dynamics of $n$ nascent RNAs are specifically determined by the instant reaction $\varnothing \xrightarrow{\rho} N$, whose stochastic evolution is characterized by

$$\frac{\mathrm{d}\bar{P}(n,t)}{\mathrm{d}t} = \rho[\bar{P}(n-1,t) - \bar{P}(n,t)], \quad \text{(S8)}$$

with initial condition being $\bar{P}(0,t) = 1$ and $\bar{P}(n,t) = 0$ for any positive integer $n$ (see Fig. S1 for a helpful illustration of the preceding arguments). Importantly, the conditional probability $P(n,t|0,t-\tau)$ coincides with $\bar{P}(n,\tau)$.

By defining generating functions $G(z,t) = \sum_n z^n P(n,t)$ and $\bar{G}(z,t) = \sum_n z^n \bar{P}(n,t)$, Eqs. (S7) and (S8) are transformed into

$$\begin{cases} \partial_t G(z,t) = \rho(z-1)G(z,t) - \rho \mathbb{1}_{[\tau,\infty)}(z-1)\bar{G}(z,\tau), \\ \partial_t \bar{G}(z,t) = \rho(z-1)\bar{G}(z,t), \end{cases}$$

which admits the solution

$$\bar{G}(z,t) = \exp[\rho(z-1)t],$$

and

$$G(z,t) = \exp[\rho(z-1)t^*], \quad \text{(S9)}$$

where $t^* = \min\{t,\tau\}$ and $\mathbb{1}_{[\tau,\infty)} = 1$ if and only if $t \in [\tau,\infty)$. Hence finally we obtain using

$$P(n,t) = \frac{1}{n!}\frac{\mathrm{d}^n G(z,t)}{\mathrm{d}z^n}\bigg|_{z=0},$$

that the solution to the delay CME of Model I is a Poisson distribution parametrized by $\rho t^*$. Interestingly, it is noted from Eq. (S9) that the distribution does not evolve any longer after time $\tau$, while it is continuously changing prior to $\tau$.

#### 1.3 Intractability of the analytical solution when delay is stochastic

Consider a modification of Model I

$$\varnothing \xrightarrow{\rho} N, \quad N \underset{\tau_\omega}{\Rightarrow} \varnothing,$$

whereby the delay $\tau_\omega$ is a random variable defined on a probability space and sampled from some distribution with random sample $\omega$.

Repeating the steps of the derivation as above, one finds difficulty in computing the contributions due to nascent RNA removal, i.e. event (B). Given the history leading to removal, as we previously argued, there are three contributions to this event and one finds that those associated with (ii) cannot be analytically resolved as we did earlier. Such a breakdown stems from the fact that the $n^*$ nascent RNAs born prior to time $t - \tau_\omega$ do not necessarily die in time $(t - \tau_\omega + \Delta t, t)$, thereby impacting the distribution at time $t$ and compromising the argument we used to arrive to Eqs. (S4) and (S8). Therefore, it is not possible to exactly solve the delay CME analytically in the presence of stochastic delay. These issues are also generally true for more complex models such as Models II and III in the main text.

### Supplementary Note 2 Exact solution for Model II: bursty transcription with delayed degradation

The model consists of the reactions

$$\varnothing \xrightarrow{\frac{\alpha b^i}{(1+b)^{i+1}}} iN, \quad N \underset{\tau}{\Rightarrow} \varnothing, \tag{S10}$$

where $i = 0, 1, 2, ...$ Nascent RNA $N$ is produced at a frequency $\alpha$ in bursts which are distributed according to a geometric distribution with mean $b$; a nascent molecule is removed from the system a time $\tau$ after its production, where $\tau$ is the time between initiation and termination of transcription.

Following the same logical steps as the derivation in Note 1, one can obtain a delay CME describing the stochastic dynamics:

$$\begin{aligned} \partial_t P(n,t) = &\sum_{m=1}^{\infty} \frac{\alpha b^m}{(1+b)^{m+1}} [P(n-m,t) - P(n,t)] \\ &+ \sum_{m=1}^{\infty} \frac{\alpha b^m}{(1+b)^{m+1}} \sum_{n'} P(n',t) \left[ P(n+m,t|n'+m,t-\tau) - P(n,t|n'+m,t-\tau) \right]. \end{aligned} \tag{S11}$$

#### 2.1 Solution of the delay master equation

This master equation can be solved using standard methods but it turns out that one can directly obtain $P(n,t)$ without even needing to write this equation. This simpler and more straightforward solution is presented next.

We are interested in the distribution of nascent RNA number $Y(t)$ to the system in Eq. (S10). Clearly, $Y(t)$ is a random variable defined on the integer support. Intuitively, $Y(t)$ is determined by two factors – the number of "packages" $I(t)$ (where a package stands for an event occurring before time $t$ such that the nascent RNA produced in these events has still not been subject to delayed degradation) and the number of nascent RNAs $X_i \sim \text{Geom}(\frac{b}{1+b})$ in each package $i$. Therefore, $Y(t)$ can be written in the form:

$$Y(t) = \sum_{i=1}^{I(t)} X_i,$$

thereby constituting a compound process. The event number $I(t)$ is determined by the system in Eq. (S1). Now we can compute the distribution of $Y(t)$ from that of $I(t)$ by using the generating-function property of a compound process. The generating function of $X_i$ is

$$G_X(z,t) = \frac{1}{1 - b(z-1)},$$

and that of package number $I(t)$ is given by Eq. (S9). Thus, the generating function of $Y(t)$ is

$$G_Y(z,t) = G[G_X(z,t),t] = \exp\left[\alpha t^* \left(\frac{1}{1-b(z-1)} - 1\right)\right],$$

which is simplifies to

$$G_Y(z,t) = \exp\left[\frac{\alpha t^* b(z-1)}{1-b(z-1)}\right]. \tag{S12}$$

#### 2.2 Conditions for the existence of delay induced zero-inflation

Here we derive conditions for the existence of the zero-inflated phenomenon described in the main text which is when the steady-state distribution is bimodal with a peak at zero and another peak at a non-zero value greater than 1. In steady-state conditions ($t \to \infty$), the generating function Eq. (S12) reduces to

$$G_Y(z) = \exp\left[\frac{\alpha\tau b(z-1)}{1-b(z-1)}\right],$$

from which we can compute the probabilities of having 0, 1 and 2 nascent RNAs at the gene locus:

$$\begin{aligned}
P(0) =& G_Y(0) = \exp\left(-\frac{\alpha b\tau}{1+b}\right), \\
P(1) =& \frac{\mathrm{d}G_Y(z)}{\mathrm{d}z}\bigg|_{z=0} = \frac{\alpha b\tau}{(1+b)^2}\exp\left(-\frac{\alpha b\tau}{1+b}\right), \\
P(2) =& \frac{1}{2!}\frac{\mathrm{d}^2G_Y(z)}{\mathrm{d}z^2}\bigg|_{z=0} = \frac{\alpha b^2\tau(2+2b+\rho\tau)}{2(1+b)^4}\exp\left(-\frac{\alpha b\tau}{1+b}\right).
\end{aligned}$$

We can define two ratios as follows

$$r_1 = \frac{P(1)}{P(0)} = \frac{\alpha b\tau}{(1+b)^2}, \qquad r_2 = \frac{P(2)}{P(1)} = \frac{b(2+2b+\alpha\tau)}{2(1+b)^2}.$$

When $r_1 < 1$ and $r_2 > 1$, the probability of having a single nascent RNA becomes a valley between $P(0)$ and $P(2)$, which establishes the sufficient and necessary condition for generating bimodality. This condition can be further simplified to read

$$2 + \frac{2}{b} < \alpha\tau < b + \frac{1}{b} + 2, \quad b > 1. \tag{S13}$$

### Supplementary Note 3 Steady state of Model III converges to that of Model II when transcription is bursty

Model III is comprised of four reactions

$$G \xrightarrow{\rho} G + N, \quad G \underset{\sigma_{\text{on}}}{\overset{\sigma_{\text{off}}}{\rightleftharpoons}} G^{\star}, \quad N \underset{\tau}{\Rightarrow} \varnothing. \tag{S14}$$

The gene can switch between active $G$ and inactive $G^{\star}$ states, transcribes nascent RNA $N$ while in the active state which subsequently is removed a time $\tau$ later. This is similar to the telegraph model for mature RNAs first studied in [2] except that since we are modelling nascent RNA, degradation is not first-order but described by a delay reaction modelling the time between initiation and termination of transcription.

#### 3.1 Derivation of the delay CME of Model III

For Model III, we would like to obtain the probabilities $P_0(n,t)$ and $P_1(n,t)$ for gene state being inactive and active, respectively. Similar to what we did in Supplementary Note 1, the two probabilities are contributed by three sets of events: (A) either a nascent RNA molecule is produced or gene state is changed in the time interval $(t, t+\Delta t)$; (B) a nascent RNA molecule is removed in the time interval $(t, t+\Delta t)$; (C) no reaction occurs in the time interval $(t, t+\Delta t)$.

In the infinitesimal limit $\Delta t \to 0$, the probabilities of event (A) simply follow the conventional propensity argument, and hence are

$$\begin{aligned} P_{01}(n, t+\Delta t; n, t) =& \sigma_{\text{off}} \Delta t P_1(n,t), \\ P_{10}(n, t+\Delta t; n, t) =& \sigma_{\text{on}} \Delta t P_0(n,t), \\ P_{11}(n, t+\Delta t; n-1, t) =& \rho \Delta t P_1(n-1,t), \end{aligned} \tag{S15}$$

where $P_{ij}(n, t+\Delta t; n', t)$ is the joint distribution of finding a cell having $n'$ and $n$ nascent RNA molecules while the gene state is in state $j$ and $i$ at time $t$ and $t+\Delta t$, respectively.

By using similar arguments in Supplementary Note 1, the probability of (i) occurring is $\rho \Delta t \delta_1(j) \times P_j(n', t-\tau)$, where $\delta_1(j) = 1$ if and only if the gene state $j = 1$ in time $(t-\tau, t-\tau+\Delta t)$. Besides, the probabilistic component (ii) for event (B) then becomes

$$P_{ij}(n, t; n'+1, t-\tau+\Delta t) = P_{ij}(n, t | n', t-\tau+\Delta t) = P_{ij}(n, t | 0, t-\tau+\Delta t). \tag{S16}$$

As such, the probability of occurring event (B) is given by the product of the probability of events (i), (ii) and (iii) and summing over all values of $n'$:

$$P_{ij}(n, t+\Delta t; n+1, t) = \rho \Delta t \sum_{n'} \delta_1(j) P_j(n', t-\tau) P_{ij}(n, t | 0, t-\tau+\Delta t) \tag{S17}$$

As per the law of total probability, we have

$$\begin{cases} P_0(n, t+\Delta t) =& P_{01}(n, t+\Delta t; n, t) + P_{01}(n, t+\Delta t; n-1, t) + P_{01}(n, t+\Delta t; n+1, t) \\ & + [P_0(n,t) - P_{10}(n, t+\Delta t; n, t) - P_{01}(n+1, t+\Delta t; n, t) - P_{01}(n-1, t+\Delta t; n, t)], \\ P_1(n, t+\Delta t) =& P_{10}(n, t+\Delta t; n, t) + P_{11}(n, t+\Delta t; n-1, t) + P_{11}(n, t+\Delta t; n+1, t) \\ & + [P_1(n,t) - P_{01}(n, t+\Delta t; n, t) - P_{11}(n+1, t+\Delta t; n, t) - P_{11}(n-1, t+\Delta t; n, t)]. \end{cases} \tag{S18}$$

Summarizing Eqs. (S15)-(S18), all are simplified to the following set of delay CMEs

$$\begin{cases} \dfrac{\mathrm{d}P_0(n,t)}{\mathrm{d}t} =& -\sigma_{\text{on}} P_0(n,t) + \sigma_{\text{off}} P_1(n,t) + \sum_{n'} \rho P_1(n', t-\tau) \left[ P_{01}(n, t | 0, t-\tau) - P_{01}(n-1, t | 0, t-\tau) \right], \\ \dfrac{\mathrm{d}P_1(n,t)}{\mathrm{d}t} =& \sigma_{\text{on}} P_0(n,t) - \sigma_{\text{off}} P_1(n,t) + \rho [P_1(n-1,t) - P_1(n,t)] + \sum_{n'} \rho P_1(n', t-\tau) \left[ P_{11}(n, t | 0, t-\tau) - P_{11}(n-1, t | 0, t-\tau) \right]. \end{cases} \tag{S19}$$

Summing over all possible $n$ for the two equations in Eq. (S19), we obtain

$$\begin{cases} \dfrac{\mathrm{d}P_0(t)}{\mathrm{d}t} = -\sigma_{\mathrm{on}}P_0(t) + \sigma_{\mathrm{off}}P_1(t), \\ \dfrac{\mathrm{d}P_1(t)}{\mathrm{d}t} = \sigma_{\mathrm{on}}P_0(t) - \sigma_{\mathrm{off}}P_1(t), \end{cases}$$

initiated with the inactive state, in which $P_0(t)$ and $P_1(t)$ are the probabilities of finding a cell at the inactivated and activated state at time $t$, and their solutions are

$$P_0(t) = \frac{\sigma_{\mathrm{off}} + \sigma_{\mathrm{on}}\mathrm{e}^{-(\sigma_{\mathrm{on}}+\sigma_{\mathrm{off}})t}}{\sigma_{\mathrm{off}} + \sigma_{\mathrm{on}}}, \quad P_1(t) = \frac{\sigma_{\mathrm{on}}[1 - \mathrm{e}^{-(\sigma_{\mathrm{on}}+\sigma_{\mathrm{off}})t}]}{\sigma_{\mathrm{off}} + \sigma_{\mathrm{on}}}.$$

Besides, it is noted that

$$\sum_{n'} P_1(n', t-\tau) = P_1(t-\tau) = \frac{\sigma_{\mathrm{on}}[1 - \mathrm{e}^{-(\sigma_{\mathrm{on}}+\sigma_{\mathrm{off}})(t-\tau)}]}{\sigma_{\mathrm{off}} + \sigma_{\mathrm{on}}} = \gamma_{t-\tau}, \tag{S20}$$

for $t \geq \tau$, and $\gamma_{t-\tau} = 0$ if $t < \tau$. The $n$ nascent RNAs of the two conditional probabilities $P_{i1}(n, t|0, t-\tau)$ for $i = 0, 1$ in Eq. (S19) are produced during time $(t-\tau, t)$, and the pertinent dynamics are that of a reaction system only composed of the three non-delayed reactions in Eq. (S14). Specifically, we have

$$\begin{cases} \dfrac{\mathrm{d}\bar{P}_0(n,t)}{\mathrm{d}t} = -\sigma_{\mathrm{on}}\bar{P}_0(n,t) + \sigma_{\mathrm{off}}\bar{P}_1(n,t), \\ \dfrac{\mathrm{d}\bar{P}_1(n,t)}{\mathrm{d}t} = \sigma_{\mathrm{on}}\bar{P}_0(n,t) - \sigma_{\mathrm{off}}\bar{P}_1(n,t) + \rho[\bar{P}_1(n-1,t) - \bar{P}_1(n,t)], \end{cases} \tag{S21}$$

initiated at $\bar{P}_0(n, 0) = 0$ for any $n$, $\bar{P}_1(n, 0) = 1$ for $n = 0$ and equal to 0 otherwise, as well as $P_{i1}(n, t|0, t-\tau) = \bar{P}_i(n, \tau)$ for any $i = 0, 1$.

Let $G_i(u, t) = \sum_n (u+1)^n P_i(n, t)$ and $\bar{G}_i(u, t) = \sum_n (u+1)^n \bar{P}_i(n, t)$. We particularly define the generating function in such a form so as to simplify notations. Then, using Eq. (S20), Eqs. (S19) and (S21) lead to the generating function equations:

$$\begin{cases} \partial_t G_0 = -\sigma_{\mathrm{on}}G_0 + \sigma_{\mathrm{off}}G_1 - \rho\gamma_{t-\tau}u\mathbb{1}_{[\tau,\infty)}\bar{G}_0^{\tau}, \\ \partial_t G_1 = \rho u G_1 + \sigma_{\mathrm{on}}G_0 - \sigma_{\mathrm{off}}G_1 - \rho\gamma_{t-\tau}u\mathbb{1}_{[\tau,\infty)}\bar{G}_1^{\tau}, \end{cases} \tag{S22}$$

and

$$\begin{cases} \partial_t \bar{G}_0 = -\sigma_{\mathrm{on}}\bar{G}_0 + \sigma_{\mathrm{off}}\bar{G}_1, \\ \partial_t \bar{G}_1 = \rho u \bar{G}_1 + \sigma_{\mathrm{on}}\bar{G}_0 - \sigma_{\mathrm{off}}\bar{G}_1, \end{cases} \tag{S23}$$

where the arguments $u$ and $t$ in the generating functions are suppressed for clarity and the superscript $\tau$ is used to emphasize the generating function $\bar{G}_i$ at the particular time $\tau$. The initial condition of Eq. (S23) is $\bar{G}_0 = 0$ and $\bar{G}_1 = 1$ when $t = 0$.

Under the condition of steady state, Eqs. (S22) reduces to

$$\begin{cases} 0 = -\sigma_{\mathrm{on}}G_0 + \sigma_{\mathrm{off}}G_1 - \rho\bar{\gamma}u\bar{G}_0^{\tau}, \\ 0 = \rho u G_1 + \sigma_{\mathrm{on}}G_0 - \sigma_{\mathrm{off}}G_1 - \rho\bar{\gamma}u\bar{G}_1^{\tau}, \end{cases} \tag{S24}$$

with $\bar{\gamma} = \sigma_{\mathrm{on}}/(\sigma_{\mathrm{on}} + \sigma_{\mathrm{off}})$.

### 3.2 Model reduction under bursty conditions

We now find an approximate solution to Model III when $\sigma_{\mathrm{off}} \gg \sigma_{\mathrm{on}}$. To this end, if we define $\delta = \sigma_{\mathrm{off}}/\sigma_{\mathrm{on}}$ and divide both sides of Eqs. (S23) by $\sigma_{\mathrm{on}}$ then we obtain

$$\begin{cases} \partial_s \bar{G}_0 = -\bar{G}_0 + \delta\bar{G}_1, \\ \partial_s \bar{G}_1 = \tilde{\rho}u\bar{G}_1 + \bar{G}_0 - \delta\bar{G}_1, \end{cases} \tag{S25}$$

with $\tilde{\rho} = \rho/\sigma_{\text{on}}$ and $s = \sigma_{\text{on}} t$. Then, we further divide all equations in Eqs. (S25) by $\delta$ and denote $\epsilon = 1/\delta$ to obtain

$$\begin{cases} \epsilon \partial_s \bar{G}_0 = -\epsilon \bar{G}_0 + \bar{G}_1, \\ \epsilon \partial_s \bar{G}_1 = bu\bar{G}_1 + \epsilon \bar{G}_0 - \bar{G}_1, \end{cases} \tag{S26}$$

with $b = \rho/\sigma_{\text{off}}$ (the mean nascent RNA burst size). From Table 1 of Ref. [3] for many genes $\epsilon$ is reported to be a small positive real number, and hence as per perturbation theory, we postulate that all the generating functions $\bar{G}_0$ and $\bar{G}_1$ have a series expansion in $\epsilon$ of the form

$$\bar{G}_0 = \bar{G}_0^{(0)} + \epsilon \bar{G}_0^{(1)} + \mathcal{O}(\epsilon^2), \qquad \bar{G}_1 = \bar{G}_1^{(0)} + \epsilon \bar{G}_1^{(1)} + \mathcal{O}(\epsilon^2).$$

Matching the coefficients of different orders in $\epsilon$ in Eq. (S26), the following set of equations ensues

$$\text{Order } \epsilon^0: \quad \bar{G}_1^{(0)} = 0,$$

$$\text{Order } \epsilon^1: \quad \begin{cases} \partial_s \bar{G}_0^{(0)} = -\bar{G}_0^{(0)} + \bar{G}_1^{(1)}, \\ \partial_s \bar{G}_1^{(0)} = bu\bar{G}_1^{(1)} + \bar{G}_0^{(0)} - \bar{G}_1^{(1)}, \end{cases}$$

all of which reduce to

$$\partial_s \bar{G}_0^{(0)} = \frac{bu}{1 - bu} \bar{G}_0^{(0)}.$$

Thus, we have

$$\bar{G}_0^{(0)} = C \exp\left(\frac{\sigma_{\text{on}} but}{1 - bu}\right),$$

where the integration constant is $C = 1/(1 - bu)$. As the process represented by Eq. (S21) is initiated at the activated gene state and quickly converges to the slow manifold (the inactivated state), the number of nascent RNAs produced during such a short ON window is subject to the geometric distribution with mean burst size $b$, whose generating function is $1/(1 - bu)$.

Directly solving Eq. (S24), we have

$$G_0 = \frac{\sigma_{\text{off}}(\bar{G}_0^\tau + \bar{G}_1^\tau) - \rho u \bar{G}_0^\tau}{\sigma_{\text{on}} + \sigma_{\text{off}}}, \qquad G_1 = \frac{\sigma_{\text{on}}(\bar{G}_0^\tau + \bar{G}_1^\tau)}{\sigma_{\text{on}} + \sigma_{\text{off}}}.$$

Furthermore, it is obtained that

$$G = G_0 + G_1 = (\bar{G}_0^\tau + \bar{G}_1^\tau) - \frac{\rho u \bar{G}_0^\tau}{\sigma_{\text{on}} + \sigma_{\text{off}}} \approx (1 - bu)\bar{G}_0^{(0)}|_{t=\tau} = \exp\left(\frac{\sigma_{\text{on}} bu\tau}{1 - bu}\right), \tag{S27}$$

where the approximation step is performed under the condition $\sigma_{\text{off}} \gg \sigma_{\text{on}}$. By comparing the generating functions in Eqs. (S12) and (S27), we find that they are the same if $\alpha = \sigma_{\text{on}}$, thereby concluding that Model III converges to Model II at steady state in bursty conditions.

### Supplementary Note 4 Neural Network Chemical Master equations for Models II and III

The Neural Network Chemical Master equation (NN-CME) for Model I was presented in the main text. Here we show the same for Models II and III.

#### 4.1 Model II

Similar to the reasoning for Model I, since only the removal of nascent RNAs is delayed, the NN-CME of the system Eq. (S10) is readily written as a term for bursty production (which is the same as in the conventional CME) plus effective degradation terms:

$$\partial_t P(n,t) = \sum_{m=1}^{\infty} \frac{\alpha b^m}{(1+b)^{m+1}}[P(n-m,t) - P(n,t)] + \mathrm{NN}_\theta(n+1,t)P(n+1,t) - \mathrm{NN}_\theta(n,t)P(n,t). \tag{S28}$$

Rewriting Eq. (S28) into the form

$$\frac{\mathrm{d}}{\mathrm{d}t}\mathbf{P}(t) = \mathbf{A}_\theta(t)\mathbf{P}(t), \tag{S29}$$

where $\mathbf{A}_\theta(t) = \mathbf{D} + \mathbf{N}_\theta(t)$, we find

$$\mathbf{D} = \begin{bmatrix} -\frac{\alpha b}{1+b} & 0 & \cdots & 0 & 0 \\ \frac{\alpha b}{(1+b)^2} & -\frac{\alpha b}{1+b} & \cdots & 0 & 0 \\ \vdots & \vdots & \ddots & \vdots & \\ \frac{\alpha b^{N-1}}{(1+b)^N} & \frac{\alpha b^{N-2}}{(1+b)^{N-1}} & \cdots & -\frac{\alpha b}{1+b} & 0 \\ \frac{\alpha b^N}{(1+b)^{N+1}} & \frac{\alpha b^{N-1}}{(1+b)^N} & \cdots & \frac{\alpha b}{(1+b)^2} & -\frac{\alpha b}{1+b} \end{bmatrix},$$

and

$$\mathbf{N}_\theta(t) = \begin{bmatrix} 0 & \mathrm{NN}_\theta(1,t) & \cdots & 0 & 0 \\ 0 & -\mathrm{NN}_\theta(1,t) & \cdots & 0 & 0 \\ \vdots & \vdots & \ddots & \vdots & \vdots \\ 0 & 0 & \cdots & -\mathrm{NN}_\theta(N-1,t) & \mathrm{NN}_\theta(N,t) \\ 0 & 0 & \cdots & 0 & -\mathrm{NN}_\theta(N,t) \end{bmatrix}.$$

#### 4.2 Model III

By similar arguments as above, the NN-CME of Model III is

$$\begin{cases} \frac{\mathrm{d}}{\mathrm{d}t}P_0(n,t) = \sigma_{\mathrm{off}}P_1(n,t) - \sigma_{\mathrm{on}}P_0(n,t) + \mathrm{NN}_{0,\theta}(n+1,t)P_0(n+1,t) - \mathrm{NN}_{0,\theta}(n,t)P_0(n,t), \\ \frac{\mathrm{d}}{\mathrm{d}t}P_1(n,t) = \sigma_{\mathrm{on}}P_0(n,t) - \sigma_{\mathrm{off}}P_1(n,t) + \mathrm{NN}_{1,\theta}(n+1,t)P_1(n+1,t) - \mathrm{NN}_{1,\theta}(n,t)P_1(n,t) + \rho[P_1(n-1,t) - P_1(n,t)]. \end{cases}$$

These equations can be rewritten into the form of Eq. (S29) where the probability vector is $\mathbf{P}(t) = [P_0(0,t),\cdots,P_0(N,t),P_1(0,t),\cdots,P_1(N,t)]^\top$, and $\mathbf{A}_\theta(t,\tau) = \mathbf{D} + \mathbf{N}_\theta(t)$ are defined as

$$\mathbf{D} = \begin{bmatrix} \mathbf{D_{11}} & \mathbf{D_{12}} \\ \mathbf{D_{21}} & \mathbf{D_{22}} \end{bmatrix},$$

and

$$\mathbf{D}_{11} = -\sigma_{\mathrm{on}}\mathbf{I}_{(N+1)\times(N+1)}, \quad \mathbf{D_{12}} = \sigma_{\mathrm{off}}\mathbf{I}_{(N+1)\times(N+1)}, \quad \mathbf{D_{21}} = \sigma_{\mathrm{on}}\mathbf{I}_{(N+1)\times(N+1)},$$

and

$$
\mathbf{D_{22}} = \begin{bmatrix} -\sigma_{\text{off}} - \rho & 0 & 0 & \cdots & 0 & 0 \\ \rho & -\sigma_{\text{off}} - \rho & 0 & \cdots & 0 & 0 \\ \vdots & \vdots & \vdots & \ddots & \vdots & \\ 0 & 0 & 0 & \cdots & -\sigma_{\text{off}} - \rho & 0 \\ 0 & 0 & 0 & \cdots & \rho & -\sigma_{\text{off}} \end{bmatrix},
$$

$$
\mathbf{N}_\theta(t) = \begin{bmatrix} \mathbf{N}_{0,\theta}(t) & 0 \\ 0 & \mathbf{N}_{1,\theta}(t) \end{bmatrix},
$$

and

$$
\mathbf{N}_{0,\theta}(t) = \begin{bmatrix} 0 & \mathrm{NN}_{0,\theta}(1,t) & 0 & \cdots & 0 & 0 \\ 0 & -\mathrm{NN}_{0,\theta}(1,t) & \mathrm{NN}_{0,\theta}(2,t) & \cdots & 0 & 0 \\ 0 & 0 & -\mathrm{NN}_{0,\theta}(2,t) & \cdots & 0 & 0 \\ \vdots & \vdots & \vdots & \ddots & \vdots & \\ 0 & 0 & 0 & \cdots & -\mathrm{NN}_{0,\theta}(N-1,t) & \mathrm{NN}_{0,\theta}(N,t) \\ 0 & 0 & 0 & \cdots & 0 & -\mathrm{NN}_{0,\theta}(N,t) \end{bmatrix},
$$

and

$$
\mathbf{N}_{1,\theta}(t) = \begin{bmatrix} 0 & \mathrm{NN}_{1,\theta}(1,t) & 0 & \cdots & 0 & 0 \\ 0 & -\mathrm{NN}_{1,\theta}(1,t) & \mathrm{NN}_{1,\theta}(2,t) & \cdots & 0 & 0 \\ 0 & 0 & -\mathrm{NN}_{1,\theta}(2,t) & \cdots & 0 & 0 \\ \vdots & \vdots & \vdots & \ddots & \vdots & \\ 0 & 0 & 0 & \cdots & -\mathrm{NN}_{1,\theta}(N-1,t) & \mathrm{NN}_{1,\theta}(N,t) \\ 0 & 0 & 0 & \cdots & 0 & -\mathrm{NN}_{1,\theta}(N,t) \end{bmatrix}.
$$

Note that the same set of matrices are used to solve Model III when the delay is stochastic (as in Fig. 4c of the main text), wherein the values of prefix biases are set to the mean of the stochastic delay ($\langle\tau\rangle$).

### Supplementary Note 5 Oscillatory gene regulatory network

Here we describe in more detail the stochastic auto-regulatory model studied in the main text and illustrated in Fig. 5a therein. The reactions comprising the model are:

$$\varnothing \xrightarrow{J_1(Y)} X, \qquad X \underset{\tau}{\Rightarrow} Y, \qquad Y \xrightarrow{J_2(Y)} \varnothing, \tag{S30}$$

with

$$J_1(Y) = k_1 S \frac{K_{\mathrm{d}}^p}{K_{\mathrm{d}}^p + Y^p}, \qquad J_2(Y) = k_2 E_{\mathrm{T}} \frac{Y}{K_{\mathrm{m}} + Y}.$$

The two reaction rates $J_1(Y)$ and $J_2(Y)$ in Eq. (S30) are in the form of a Hill function. The synthesis rate $J_1(Y)$ of premature protein $X$ is proportional to the number of upstream regulators $S$, and is downregulated by the number of protein $Y$. The dissociation constant for protein Y and the gene promoter is $K_{\mathrm{d}}$, and $p$ is the Hill coefficient. In the degradation rate $J_2(Y)$, $E_{\mathrm{T}}$ stands for the total number of protease, $k_2$ is the turnover rate, and $K_{\mathrm{m}}$ is the Michaelis constant.

Let $P(x, y, t)$ be the probability of finding $x$ molecules of premature protein $X$ and $y$ molecules of protein $Y$ in a cell at time $t$. Then by similar arguments used for previous models, the NN-CME governing the time-evolution of the probability $P(x, y, t)$ is given by

$$\frac{\mathrm{d}}{\mathrm{d}t} P(x,y,t) = (\mathbb{E}^{-1,0} - 1) J_1(y) P(x,y,t) + (\mathbb{E}^{0,1} - 1) J_2(y) P(x,y,t) + (\mathbb{E}^{1,-1} - 1) \mathrm{NN}_\theta(x,y,t) P(x,y,t), \tag{S31}$$

with $\mathbb{E}$ being the step operator whose function is $\mathbb{E}^{i,j} f(x,y) = f(x+i, y+j)$ and initial condition $P(x, y, 0) = 1$ if and only if $x = y = 0$.

Again we are able to rewrite Eq. (S31) in the compact form Eq. (S29) by selecting a sufficiently large $N$ such that $P(x, y, t) = 0$ for any $x \geq N$ and $y \geq N$ and stacking $P(x, y, t)$ to form a vector of length $(N+1)^2$:

$$\mathbf{P}(t) = [P(0,0,t), \ \cdots, \ P(N,0,t), P(0,1,t), \ \cdots, \ P(N,1,t), \ \cdots, P(0,N,t), \ \cdots, \ P(N,N,t)]^\top.$$

Besides, the following matrices are needed to assemble matrix $\mathbf{A}_\theta(t) = \mathbf{D}_1 + \mathbf{D}_2 + \mathbf{N}_\theta(t)$, and $\mathbf{D}_1 = \mathrm{diag}\{\mathbf{D}_1^0, \cdots, \mathbf{D}_1^N\} \in \mathbb{R}^{(N+1)^2 \times (N+1)^2}$ with

$$\mathbf{D}_1^y = \begin{bmatrix} -J_1(y) & 0 & \cdots & 0 & 0 \\ J_1(y) & -J_1(y) & \cdots & 0 & 0 \\ \vdots & \vdots & \vdots & \ddots & \vdots \\ 0 & 0 & \cdots & -J_1(y) & 0 \\ 0 & 0 & \cdots & J_1(y) & 0 \end{bmatrix} \in \mathbb{R}^{(N+1)\times(N+1)},$$

and

$$\mathbf{D}_2 = \begin{bmatrix} -\mathbf{D}_2^0 & \mathbf{D}_2^1 & \cdots & 0 & 0 \\ 0 & -\mathbf{D}_2^1 & \cdots & 0 & 0 \\ \vdots & \vdots & \vdots & \ddots & \vdots \\ 0 & 0 & \cdots & -\mathbf{D}_2^{N-1} & \mathbf{D}_2^N \\ 0 & 0 & \cdots & 0 & -\mathbf{D}_2^N \end{bmatrix} \in \mathbb{R}^{(N+1)^2\times(N+1)^2}$$

and

$$\mathbf{D}_2^y = J_2(y) \mathbf{I}_{(N+1)\times(N+1)}$$

and

$$\mathbf{N}_\theta(t) = \begin{bmatrix} -\mathbf{N}_\theta^{a,0} & 0 & \cdots & 0 & 0 \\ \mathbf{N}_\theta^{b,0} & -\mathbf{N}_\theta^{a,1} & \cdots & 0 & 0 \\ \vdots & \vdots & \vdots & \ddots & \vdots \\ 0 & 0 & \cdots & -\mathbf{N}_\theta^{a,N-1} & 0 \\ 0 & 0 & \cdots & \mathbf{N}_\theta^{b,N-1} & 0 \end{bmatrix} \in \mathbb{R}^{(N+1)^2\times(N+1)^2}$$

and

$$\mathbf{N}_\theta^{a,k} = \mathrm{diag}\{\mathrm{NN}_\theta(0,k,t),\cdots,\mathrm{NN}_\theta(N-1,k,t),0\} \in \mathbb{R}^{(N+1)\times(N+1)},$$

for $k = 1,\cdots,N$, and

$$\mathbf{N}_\theta^{b,k} = \begin{bmatrix} 0 & \mathrm{NN}_\theta(1,k,t) & \cdots & 0 & 0 & 0 \\ 0 & 0 & \cdots & 0 & 0 & 0 \\ \vdots & \vdots & \vdots & \ddots & \vdots & \vdots \\ 0 & 0 & \cdots & 0 & \mathrm{NN}_\theta(N,k-1,t) & 0 \\ 0 & 0 & \cdots & 0 & 0 & 0 \\ 0 & 0 & \cdots & 0 & 0 & 0 \end{bmatrix} \in \mathbb{R}^{(N+1)\times(N+1)},$$

for $k = 0,\cdots,N-1$. The preset biases $r_n$ of $\mathrm{NN}_\theta(n,k,t)$ are increasing functions of $n$ and proportional to $\tau^{-1}$.

### Supplementary Note 6 Calculation of confidence intervals using profile likelihood

The confidence intervals are derived by the concept of profile likelihood and the nuisance parameters (see p. 264-265 in Ref. [4]).

With reference to Fig. 6 of the main text, suppose that the burst frequency $\alpha$ is the parameter that we are interested in estimating its confidence interval. The augmented parameter vector is $\psi = [\alpha, b, \theta^\top]^\top$, and the nuisance parameters accordingly become $\lambda = [b, \theta^\top]^\top$. Let $(\hat{\alpha}, \hat{\lambda})$ be the solution minimizing the mean squared error cost function to the NN-CME, and potentially $\hat{\psi}$ minimize the negative log-likelihood:

$$(\hat{\alpha}, \hat{\lambda}) = \arg\min_{(\alpha,\lambda)} \mathcal{L}(\alpha, \lambda),$$

where

$$\mathcal{L}(\alpha, \lambda) = -\sum_{i,j} \ln P_\psi(x_{i,t_j}),$$

and $P_\psi$ is the probability of observing $x_{i,t_j}$ molecules at time $t_j$, which is predicted by NN-CME.

As the burst frequency $\alpha$ is the one we would like to quantify uncertainty, we define a new type of optimization problem:

$$\hat{\lambda}_\alpha = \arg\min_\lambda \mathcal{L}(\alpha, \lambda),$$

in which it means we attempt to minimize the likelihood by only tuning $\lambda$ and fixing the value $\alpha$.

Hence, we are able to define a function of $\alpha$ as

$$\Delta\mathcal{L}(\alpha) = \mathcal{L}(\alpha, \hat{\lambda}_\alpha) - \mathcal{L}(\hat{\alpha}, \hat{\lambda}).$$

Asymptotic theory [4] shows that

$$\Delta\mathcal{L}(\alpha) \xrightarrow{d} \frac{1}{2}\chi_1^2.$$

Thus, by varying $\alpha$, we are able to obtain a curve for $\Delta\mathcal{L}$ (see the right panel of Fig. 6a main text), whose intersections with the horizontal line 1.92 give the 95% confidence interval. The values of $\hat{\lambda}_\alpha$ can be quickly computed by retraining NN-CME initialized with $\hat{\psi}$. The 95% confidence intervals for burst size $b$ or Model III can be obtained similarly.

**Neural network specifications:**

The following table summarizes the technical specifications about the ANN used for all the examples presented in the main text. Note that there are two hyperparameters involved in the NN-CME, namely the number of hidden neurons and the learning rate. We found that the results are not very sensitive to the number of hidden neurons, and below we just present a set of hyperparameters that works and achieved good results. Second, the training can be started with a smaller learning rate to achieve an acceptable fitting performance, and then the learning rate can be increased to achieve faster convergence. This is the reason why a step-wise setting is used in the experiments.

Table 1: **Technical specifications about the neural networks.** The details about the structure of neural networks, training, SSA data used for training and model are included. The decay profile of learning rate $\ell$ labeled with $^\dagger$ is presented in the form that $\sum_i$ learning rate $i \times$ epochs $i$, which means a neural network is trained for epochs $i$ times with the learning rate $i$.

| Fig. | Neural network structure | | Training | SSA data | | Model | | | | | | | | | | | |
|---|---|---|---|---|---|---|---|---|---|---|---|---|---|---|---|---|---|
| | Truncation $N$ | Hidden neurons # | Learning rate $\ell^\dagger$ | Snapshot # ($N_{\text{shot}}$) | Sample # ($N_{\text{sim}}$) | Eq. | $\rho$ | $\tau$ | $\sigma_{\text{on}}$ | $\sigma_{\text{off}}$ | $\alpha$ | $b$ | $k_1 \times S$ | $K_{\text{d}}$ | $p$ | $k_2 \times E_{\text{T}}$ | $K_{\text{m}}$ |
| 2b-1 | 296 | 100 | $0.01 \times 50$ | 20 | $10^3$ | (S1) | 20 | 10 | / | / | / | / | / | / | / | / | / |
| 2b-2 | 70 | 10 | $0.01 \times 30 + 0.005 \times 70$ | 100 | $2 \times 10^4$ | (S10) | / | 130 | / | / | 0.0282 | 3.46 | / | / | / | / | / |
| 2b-3 | 118 | 300 | $0.01 \times 20 + 0.005 \times 30$ | 40 | $8 \times 10^3$ | (S14) | 2.11 | 200 | 0.0282 | 0.609 | / | / | / | / | / | / | / |
| 2b-4 | 45 | 500 | $0.01 \times 20 + 0.005 \times 30$ | 100 | $10^4$ | (S14) | 20 | 1 | 1 | 1 | / | / | / | / | / | / | / |
| 3 | 296 | 200 | $0.25 \times 7 + 0.12 \times 8 + 0.02 \times 2$ | 3/6/9 | $0.1/0.3/1/3/10/30 \times 10^3$ | (S1) | 20 | 10 | / | / | / | / | / | / | / | / | / |
| 4c-2 | 108 | 300 | $0.01 \times 20 + 0.005 \times 30$ | 40 | $10^4$ | (S14) | 2.11 | $\text{logn}(1, \sqrt{2}) + 120$ | 0.0282 | 0.609 | / | / | / | / | / | / | / |
| 4c-3 | 107 | 300 | $0.01 \times 20 + 0.005 \times 30$ | 40 | $10^4$ | (S14) | 2.11 | $\text{logn}(0, 2) + 120$ | 0.0282 | 0.609 | / | / | / | / | / | / | / |
| 5 | 25 | 100 | $0.01 \times 20 + 0.001 \times 50$ | 50 | $1.5 \times 10^4$ | (S30) | / | 10 | / | / | / | / | 1 | 1 | 2 | 1 | 1 |
| 6-1 | 20 | 20 | $0.01 \times 100 + 0.005 \times 200$ | 100 | $10^4$ | (S10) | / | 388 | / | / | 0.0011 | 1.12 | / | / | / | / | / |
| 6-2 | 20 | 20 | $0.01 \times 100 + 0.005 \times 200$ | 100 | $10^4$ | (S10) | / | 60 | / | / | 0.0072 | 1.95 | / | / | / | / | / |
| 6-3 | 20 | 20 | $0.01 \times 100 + 0.005 \times 200$ | 100 | $10^4$ | (S10) | / | 578 | / | / | 0.0038 | 1.38 | / | / | / | / | / |
| 6-4 | 20 | 20 | $0.01 \times 100 + 0.005 \times 200$ | 100 | $10^4$ | (S10) | / | 1068 | / | / | 0.0016 | 1.39 | / | / | / | / | / |
| 6-5 | 20 | 20 | $0.01 \times 100 + 0.005 \times 200$ | 100 | $10^4$ | (S10) | / | 1081 | / | / | 0.0008 | 2.11 | / | / | / | / | / |
| S2 | 35 | 100 | $0.01 \times 100 + 0.005 \times 100$ | 1 | $0.1/0.3/1/3/10/30 \times 10^3$ | (S10) | / | 578 | / | / | 0.0038 | 1.38 | / | / | / | / | / |
| S3-1 | 30 | 5 | $0.01 \times 50 + 0.005 \times 200 + 0.001 \times 100$ | 100 | $1.5 \times 10^4$ | (S14) | 0.077 | 103 | 0.0029 | 0.002 | / | / | / | / | / | / | / |
| S3-2 | 30 | 10 | $0.01 \times 50 + 0.005 \times 200 + 0.001 \times 100$ | 100 | $1.5 \times 10^4$ | (S14) | 0.139 | 32 | 0.0043 | 0.015 | / | / | / | / | / | / | / |

| Fig. | Sample collected time |
|---|---|
| 2b-1 | $\{0, 2, 4, \cdots, 40\}$ |
| 2b-2 | $\{0, 2, 4, \cdots, 200\}$ |
| 2b-3 | $\{0, 20, 40, \cdots, 800\}$ |
| 2b-4 | $\{0, 0.1, 0.2, \cdots, 10\}$ |
| 3 | $\{4, 8, 28\}$, $\{4, 8, 14, 26, 38, 50\}$, $\{4, 8, 14, 20, 26, 32, 38, 44, 50\}$ |
| 4c-2 | $\{0, 50, 100, \cdots, 2000\}$ |
| 4c-3 | $\{0, 50, 100, \cdots, 2000\}$ |
| 5 | $\{0, 2, 4, \cdots, 100\}$ |
| 6c-1 | $\{0, 40, 80, \cdots, 4000\}$ |
| 6c-2 | $\{0, 6, 12, \cdots, 600\}$ |
| 6c-3 | $\{0, 50, 100, \cdots, 5000\}$ |
| 6c-4 | $\{0, 100, 200, \cdots, 10000\}$ |
| 6c-5 | $\{0, 100, 200, \cdots, 10000\}$ |
| S3-1 | $\{0, 10, 20, \cdots, 1000\}$ |
| S3-2 | $\{0, 10, 20, \cdots, 1000\}$ |

**Supplementary Figures**

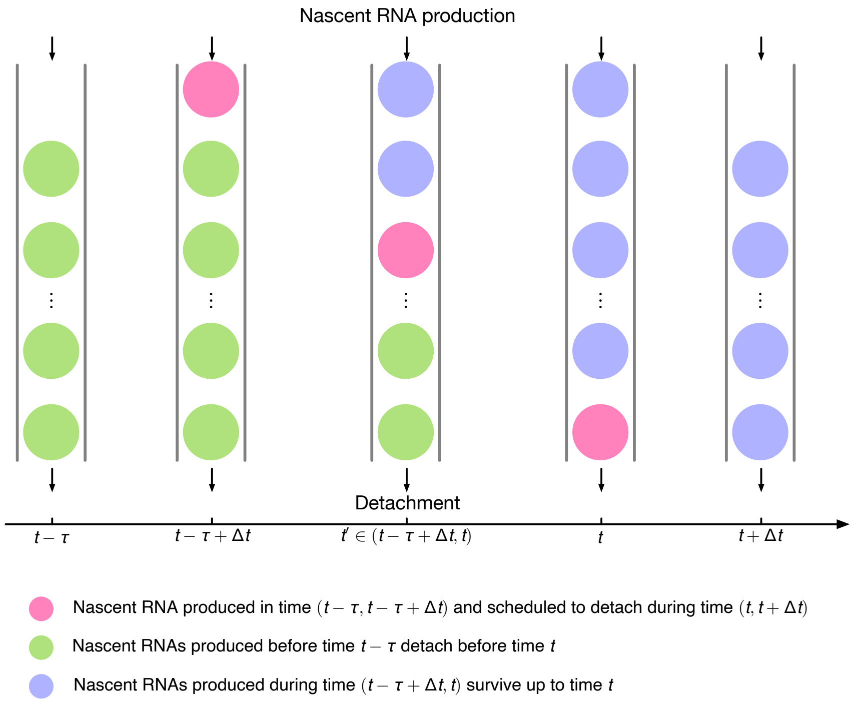

Figure S1: Illustration of the chain of events culminating in the removal (detachment) of a nascent RNA in Model I (refer to event (B) in Note I). Let's assume there are $n'$ nascent RNAs (green balls) produced before time $t-\tau$. In the next time step $t-\tau+\Delta t$, there is a new born RNA (red ball). During $(t-\tau+\Delta t, t)$, the first $n'$ green RNAs are progressively removed and in the meanwhile $n$ new RNAs (blue balls) are produced. Since each nascent RNA stays for a fixed time $\tau$, the red RNA definitely leaves in the time interval $(t, t+\Delta t)$. Note that during the time interval $(t-\tau+\Delta t, t)$, the red RNA does not participate in any dynamics of the system which explains Eq. (S3) and the dynamics of the blue RNAs follows Eq. (S8) which is a simple birth process from $(0, \tau)$.

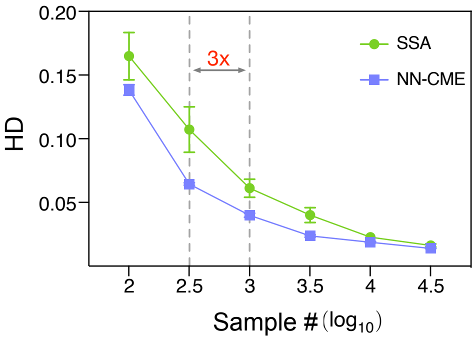

Figure S2: Testing the precision and computation efficiency of the ANN-aided model approximation for Model II using a steady-state training procedure, i.e. we solve the algebraic equations $\mathbf{A}_\theta(t)\mathbf{P}(t) = 0$ during training. Note that the steady-state training only uses one snapshot of histogram. It shows the Hellinger distance (HD) between the distribution of the NN-CME and histograms computed from stochastic simulations (SSA) as a function of sample size. Each SSA data point is averaged over 10 independent runs, while each NN-CME point is averaged over 3 independent trainings. It shows that the steady-state ANN-aided approximation is able to achieve comparable precision to that of the SSA while only using about 1/3 of the samples necessary for the latter. The kinetic parameters are: $\alpha = 3.84 \times 10^{-3}$, $b = 1.38$.

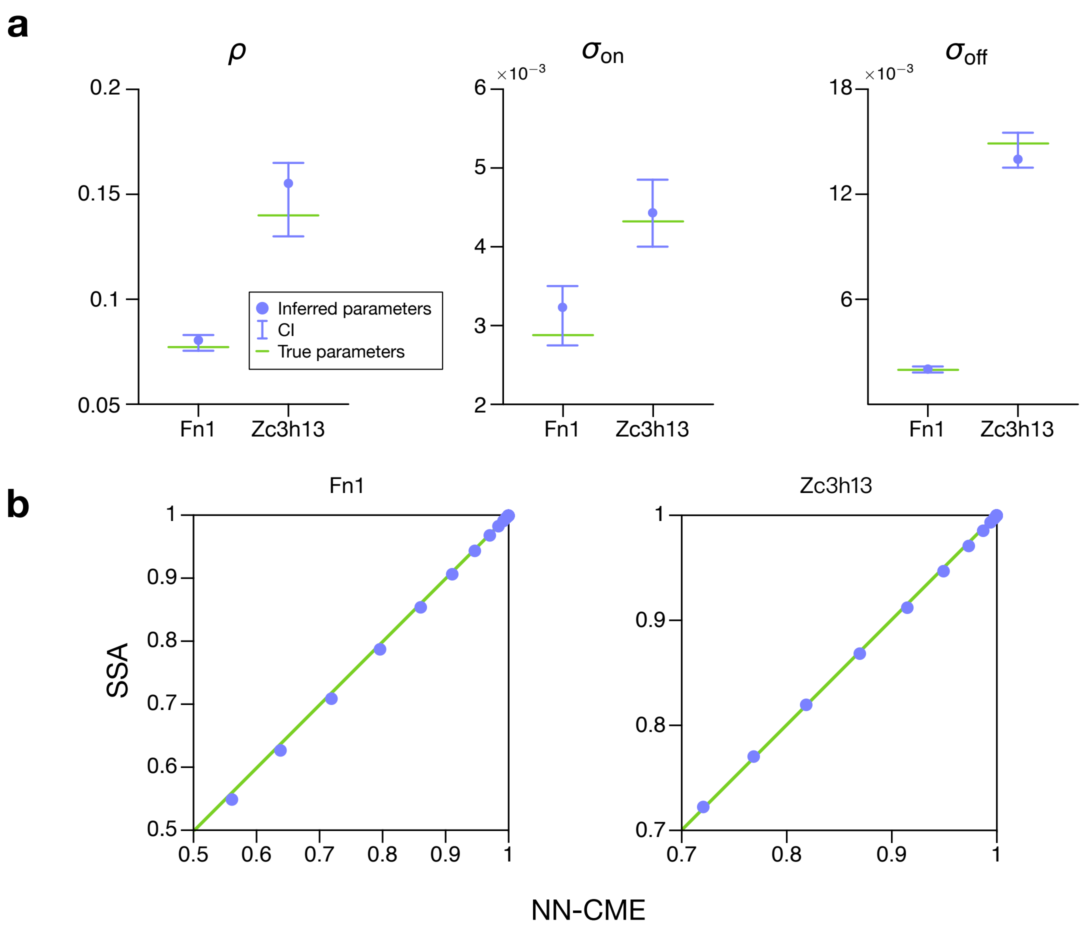

Figure S3: Inference of kinetic parameters ($\rho$, $\sigma_{\text{on}}$ and $\sigma_{\text{off}}$) for Model III using ANN-aided model approximation. (a) Shows the inferred values and the corresponding 95% confidence interval computed using the profile likelihood method (Supplementary Note 6). (b) Shows by means of quintile-quintile plots, that simultaneously with inference, the NN-CME given by the method has a steady-state distribution that is in excellent agreement with that of SSA. See Table 1 for parameters.
